# Whole-genome profiling of native 5-hydroxymethylation in human neurons with long-read sequencing

**DOI:** 10.64898/2026.01.11.695537

**Authors:** Dennis Klose, Mohammad Hossein Sepehri, Remi-André Olsen, Ha Vu, Jason Ernst, Lara Kular, Maria Needhamsen, Maja Jagodic

## Abstract

The 5-hydroxymethylcytosine (5hmC) modification of DNA is particularly prevalent in neurons and thereby a hallmark of the brain’s epigenetic landscape. While 5mC DNA methylation is a well-known player in genome stability and transcriptional regulation, the role of 5hmC remains largely unknown. Here, we used long-read Oxford Nanopore Technology (ONT) to profile whole-genome, native 5mC and 5hmC levels in sorted neuronal nuclei samples from human post-mortem brain tissue. We applied different models for DNA modification calling and compared with array-based 5mC and 5hmC levels derived from the same samples, demonstrating high sample-wise correlations. Annotation across genomic and regulatory features, as well as chromatin states, generated by the International Human Epigenome Consortium, revealed high levels of 5hmC in introns, actively transcribed genes and (distal) enhancers. Pathway analysis of genes with high levels of 5hmC (> 60%) were enriched in neuron-related terms, with functional variety when stratifying across chromatin states. Analysis of transcription factor motifs in highly methylated regions, demonstrated 5hmC- and 5mC-specific enrichment affecting downstream regulatory networks.

Altogether, our study demonstrates the potential of ONT to characterize whole-genome, native 5hmC and 5mC DNA modifications in human neurons, specifically highlighting the enrichment of 5hmC in actively transcribed regions and enhancers in the human brain.

## Introduction

The 5-methylcytosine (5mC) is the most studied epigenetic modification originally described in mammals in 1948 and since then characterized to be involved in regulating gene expression, guarding genome stability and orchestrating development [1, 2]. The process of active oxidation-dependent demethylation mediated by ten-eleven translocation (TET) enzymes was described in 2009 [3]. This process involves the conversion of 5mC to 5-hydroxymethylcytosine (5hmC), 5-formylcytosine (5fC) and 5-carboxylcytosine (5caC), finally resulting in a non-methylated cytosine. In addition to being an intermediate mark during active demethylation, 5hmC was found to be particularly enriched in neurons implicating its functional role [4–7]. While the regulatory mechanisms are not yet clearly understood, levels of 5hmC in gene bodies, chromatin enhancer states and transcriptional start sites (TSS) have been found to be associated with gene expression in a tissue-specific manner [3, 8, 9].

Traditionally, 5mC levels have been measured by methods such as whole genome bisulfite sequencing (WGBS), reduced representation bisulfite sequencing (RRBS) or Illumina BeadChips DNA methylation arrays (27K, 450K, 850K) following treatment with sodium bisulfite (BS) [10]. While broadly used and generally reliable, those methods encompass different biases and shortcomings, such as (i) degradation of DNA, due to the stern chemical treatment of the genomic DNA with BS for conversion of non-methylated cytosines, (ii) complexity reduction of the genome to three nucleotides, and (iii) inability to cover a large representative amount of CpG positions throughout the entire genome [11]. Importantly, BS conversion does not enable discrimination between true 5mC and 5hmC resulting in all these methods quantifying a net effect of the two modifications. The detection of 5hmC typically requires another parallel treatment of a separate aliquot of the sample either with an oxidative chemical reagent (oxBS) or an enzymatic reaction referred to as Tet-assisted bisulfite (TAB) [12]. 5hmC levels are subsequently calculated indirectly, for example through subtraction of oxidative BS (true 5mC) signals from total BS [13]. Thus, the accurate estimation of true 5mC and 5hmC modifications remains insufficiently resolved.

The possibility to detect native 5mC and 5hmC modifications simultaneously from the same sample on a genome-wide level with Oxford Nanopore Technology (ONT) opens up a new approach to epigenetic analyses [14]. ONT technology omits BS and enzymatic conversion steps and profiles native genomic DNA without potential amplification bias as commonly introduced in traditional methods. Rather, nucleotide modifications are detected as changes in electric currents when long, native genomic DNA fragments pass through pores [15]. Base-and modification calling algorithms, trained on existing genetic and epigenetic data using deep learning, were developed together with the technology [16].

In this study, we aimed to explore the potential of ONT for profiling of 5hmC in human neurons by comparing different models for DNA modification calling and benchmarking against the array-based TrueMethyl BS/oxBS methodology, applied to the same samples. We annotated 5hmC and 5mC levels based on common regulatory and genomic features, brain-related chromatin states and furthermore conducted pathway and transcription factor enrichment analysis, giving insights into unique characteristics of regulatory features in the brain.

## Methods

### Sample description and data accessibility

Two post-mortem neuronal samples were obtained from the Multiple Sclerosis and Parkinson’s Tissue Bank (Imperial College London). As previously described [17], the NeuN^+^ nuclei were isolated using fluorescence activated nuclei sorting (FANS) from one pathology-free Normal Appearing White Matter (NAWM) brain specimen of a person with Multiple Sclerosis (MS) (S1) and one white matter sample from a non-neurological control (NNC) (S2) (Table 1). 5mC and 5hmC values had previously been obtained from the same two samples, which had been processed in parallel with bisulfite (BS)- and oxidative BS-based methods followed by Illumina Infinium Human450KBeadChip (450K) profiling [17] with data available at the Gene Expression Omnibus (GEO) database under GSE119532. Newly generated ONT data can be accessed at the European Genome-phenome Archive (EGA) under the (to be added) accession number.

**Table 1.** Description of the post-mortem neuronal samples.

| Group | Sample | Sex | Age | ONT | Illumina 450K |  |
| --- | --- | --- | --- | --- | --- | --- |
|  |  |  |  | ID | Array | BS, oxBS |
| MS (S1) | MS076-A1D3 | F | 49 | P21512_102 | 200090270050 | R04C01, R04C02 |
| NNC (S2) | PDC029-P3D3 | M | 82 | P21512_101 | 200090270050 | R06C01, R06C02 |

MS, multiple sclerosis; NNC, non-neurological control; ONT, Oxford Nanopore Technology;
BS, bisulfite Illumina array; oxBS, oxidative bisulfite Illumina array.

### Library preparation and sequencing

Purified genomic DNA was used as input for ONT library preparations with a total amount of 306 ng (fragment size: 10 322 bp) and 148 ng (fragment size: 9 343 bp) added for S1 and S2 samples, respectively. Libraries were prepared with the Genomic DNA by Ligation (SQK-LSK109) Nanopore protocol following manufactory’s guidelines. Library concentrations were measured with Qubit and a total amount of 60 ng (9.406 fmol) and 43.68 ng (7.565 fmol) were added to the R9.4.1 PromethION flow cell (FLO-PRO002). Subsequently, the following run settings were applied: -165mV Initial Bias Voltage, 90min Mux scan period, high-accuracy basecalling and min_qscore=7 read filter using the listed tools: MinKNOW (v.20.06.18), MinKNOW core (v.4.0.5), Bream (v.6.0.10) and Guppy (v.4.0.11). A runtime of 72 h generated 5.92 M reads (27.8 Gb passed and 4.3Gb failed bases) and 0.240 M reads (0.840 Gb passed and 0.428 Gb failed bases) for S1 and S2, respectively.

### Read mapping, base calling, DNA methylation calling

For base calling we used Guppy v.5.0.16 (https://github.com/nanoporetech/pyguppyclient) with the default high-accuracy (HAC) model (dna_r9.4.1_450bps_hac.cfg) through Megalodon version 2.5.0 (https://github.com/nanoporetech/megalodon) with .fast5 files as input. DNA methylation (5mC and 5hmC) calling in CpG context was conducted using the Remora (https://github.com/nanoporetech/remora) HAC model: dna_r9.4.1_e8 hac 0.0.0 5hmc_5mc CG, whereas DNA methylation calling of 5mC in all contexts, referred to as CpX, was conducted using the following config file: res_dna_r941_prom_modbases_5mC_v001.cfg. Subsequently, .fastq files were aligned to the GCA_000001405.29_GRCh38.p14 human reference genome (NCBI) with minimap2 v.2.24 [18]. A conversion table (https://genome.ucsc.edu/cgi-bin/hgTracks?db=hg38&chromInfoPage=) was used to convert alias chromosome names to “chr1”, “chr2” etc. and the coverage2cytosine function from Bismark v.0.24.2 [19] to distinguish CpG and CpH (= CHG and CHH) sites.

### Comparison of Illumina BeadChip 450K and Oxford Nanopore Technology

To compare 5mC and 5hmC levels from Illumina 450K arrays with methylation signals from ONT, we converted their genomic coordinates from hg19 to hg38. For this, we used the R packages IlluminaHumanMethylation450kanno.ilmn12.hg19 (v0.6.1), rtracklayer (v1.60.1) and GenomicRanges (v1.52.1). We joined the data on genomic position (chromosome, start, end, strand) and used two coverage cutoffs (5X and 10X) in further analysis.

### Annotation of 5hmC and 5mC marks

Reads with less than either 5X or 10X coverage and those mapping to the X- and Y-chromosome were excluded. SeqMonk (v1.48.1) (https://www.bioinformatics.babraham.ac.uk/projects/seqmonk/https://www.bioinformatics.babraham.ac.uk/projects/seqmonk/) was used to estimate DNA methylation in 3 kb running bins, within genomic (upstream, 5’UTR exon, coding exon, first intron, internal intron, last intron, 3’UTR exon and downstream) and regulatory (distal enhancers, proximal enhancers, CpG islands, promoters, exons and introns) features using the “Define Probes” in the “Bisulfite methylation over features” tool, selecting “Combined value to report” = mean. DNA methylation levels of chromatin states, including the universal/stacked 100-state chromHMM and summary 18-state chromHMM from healthy brain samples, were extracted in the same way. The chromHMM state annotations were obtained from the International Human Epigenome Consortium (IHEC), whereas the annotation for hg38 genomic and basic regulatory features was downloaded from UCSC (last updated: 20^th^ May 2020) and converted to .bed format using the bigBedToBed tool provided by UCSC for the command line (https://genome.ucsc.edu/cgi-bin/hgTables?db=hg38&hgta_group=regulation&hgta_track=encodeCcreCombined&hgta_table=encodeCcreCombined&hgta_doSchema=describe+table+schemahttps://genome.ucsc.edu/cgi-bin/hgTables?db=hg38&hgta_group=regulation&hgta_track=encodeCcreCombined&hgta_table=encodeCcreCombined&hgta_doSchema=describe+table+schema, http://hgdownload.soe.ucsc.edu/admin/exe/http://hgdownload.soe.ucsc.edu/admin/exe/). DNA methylation estimates in each defined probe were exported using the “Annotated probe report” tool. DNA methylation levels reported per cytosine base were quantified in R, except for gene body methylation levels, where the command line program CpGtools was used [20]. For CpGtools annotation we used the proposed hg38.RefSeq.union.bed file from https://sourceforge.net/projects/rseqc/files/BED/Human_Homo_sapiens_merge_transcripts/https://sourceforge.net/projects/rseqc/files/BED/Human_Homo_sapiens_merge_transcripts/. Closest genes within ± 2 kb of a chromatin state region (incl. overlapping) were determined using the “Annotate probe report” tool from SeqMonk. Annotation was performed for any 200 bp chromatin state window as provided by IHEC.

### Ingenuity Pathway Analysis

To annotate genes for Ingenuity Pathways analysis (IPA), the gencode.v38.annotation.gtf file was merged with DNA methylation sites (filtered for 10X coverage) using the GenomicRanges v. 1.60.0 and GenomicFeatures v. 3.21 Bioconductor packages [21]. Sites 1.5kb upstream of the TSS were considered to include putative promoter regions. Chromatin states, based on the healthy brain summary chromHMM annotation from IHEC was added as well. Promoters were defined as regions 1.5kb upstream of the TSS and exon 1, which overlapped with a “_Tss” state including: 1_TssA, 2_TssFlnk, 3_TssFlnkU, 4_TssFlnkD, 14_TssBiv and 15_EnhBiv. Transcribed regions (Tx) were defined as exons (from gencode.v38.annotation.gtf) overlapping with “Tx” states 5_Tx and 6_ TxWk. Enhancers were defined as regions overlapping with a “_Enh” state, including: 8_EnhG2, 9_EnhA1, 10_EnhA2 and 11_EnhWk. Repressed polycomb (PC) states were defined as regions overlapping with 16_ReprPC or 17_ReprPCWk. Zinc finger proteins (ZNF) were defined as regions overlapping with the 12_ZNF/Rpts chromatin state, which was subsequently filtered for ZNF annotated genes. Sites with 5hmC > 60% were filtered prior to IPA.

### Transcription factor motif enrichment analysis

The CentriMo tool [22], available through the MEME suite [23], was utilized for transcription factor (TF) motif enrichment analysis. 5hmC and 5mC sites above 60% and 80% DNA methylation, respectively, were selected and extended 15bp up- and down-stream of their target CpG. Control CpGs with 0% DNA methylation were utilized for relative enrichment. The Jaspar core (2024) vertebrates, non-redundant DNA motif database was selected [24]. Result output files were filtered for CpG consensus, and TFs with adj.Pval < 0.01 were considered significant.

### Transcriptional Regulatory Network Analysis

Protein-coding target genes of significantly enriched transcription factors were predicted using JASPAR 2024 (v0.99.7) motifs, based on position weight matrices (PWM) [24], which were scanned in a ±15 bp window around each CpG using TFBSTools (v1.46.0) [25] using the BSgenome.Hsapiens.UCSC.hg38 as reference (v1.4.5) [26]. Target gene expression was inferred from gene body 5hmC levels (excluding exon 1), with a threshold of >60% defined as active transcription. Transcriptional regulatory networks were rendered with visNetwork (v2.1.4) [27] and visualized in cytoScape [28] with TFs as nodes and edge colors representing enriched or depleted TF target genes estimated with Fisher’s exact test. Functional interpretation was assessed with GO enrichment (clusterProfiler (v4.16.0) [29] and per-TF enrichment bobble plots were generated with ggplot2 v4.0.0. Integration with the Human Protein Atlas single-nuclei brain data [30] allowed visualization of TF and target gene expression levels across brain cell types. Different parameters, including PWM, window size and 5hmC thresholds were assessed using the 5hmC_regulator shinyApp, which, apart from JASPAR 2024 (v0.99.7), included several additional resources such as DoRothEA regulons (v1.20.0) [31], CoLlecTRI via decoupleR (v2.140) [32, 33], TRRUST v2 [34], and allows for upload of user-supplied TF-target tables. For interpretation GO, Reactome (clusterProfiler (v4.16.0) [29] and ReactomePA (v1.52.0) [35]) were included. Access to the 5hmC_regulator shinyApp can be found here: to be added.

### Statistical analysis

Statistical analyses were performed in Rstudio (v2023.06.1+524) using R language (v4.3.1), unless stated otherwise. For plotting, we used the ggplot (v3.4.3), ggpubr (v0.6.0), ggVennDiagram (v1.5.4) [36] and ComplexHeatmap (v2.24.1) [37] packages, for general data manipulations we used dplyr (v1.1.3), tidyr (v1.3.0) and stringr (v1.5.0) [38]. For statistics we used the base R functions cor.test, lm and poly, such as rstatix (v0.7.2) functions wilcox_test and wilcox_effsize. For creating heatmaps we used the pheatmap package (v1.0.12) with the complete linkage clustering method. For figure assembly we used Affinity Designer 2.

## Results

### Oxford Nanopore technology generates long reads and whole-genome coverage of the neuronal methylome

We profiled 5mC and 5hmC in neuronal DNA isolated from NeuN^+^ nuclei sorted from post-mortem brain tissue of one MS (S1) and one NNC (S2) sample using Oxford Nanopore technology after DNA extraction, fragmentation and library preparation (Fig. 1a). Squiggle signals were generated by DNA passing through a membrane protein (“nanopore”) and transformed into base calls by Guppy. Remora was used for calling epigenetically modified bases, 5hmC and 5mC, in CpGs (cytosine followed by guanine), whereas a combined signal of the two modifications was called in all contexts (CpX). After base and DNA methylation calling, reads passing quality control were mapped to human genome hg38 using minimap2 (Fig. 1b). Since the input amount (306 ng vs. 148 ng), fragment sizes (10 322 bp vs. 9 343 bp) and number of reads (5.92M vs. 0.240M) differed between S1 and S2, respectively, we first explored sample-specific outcomes such as read length and coverage. The S1 sample yielded more and longer reads (Fig. 1c), consequently increasing the coverage in both CpG and CpX contexts (Fig. 1d). Nevertheless, both samples reached near whole-genome 1X coverage of 57.7M and 48.9M sites in the CpG context, considering both strands, for S1 and S2, respectively (Fig. 1d).

**Figure 1.**
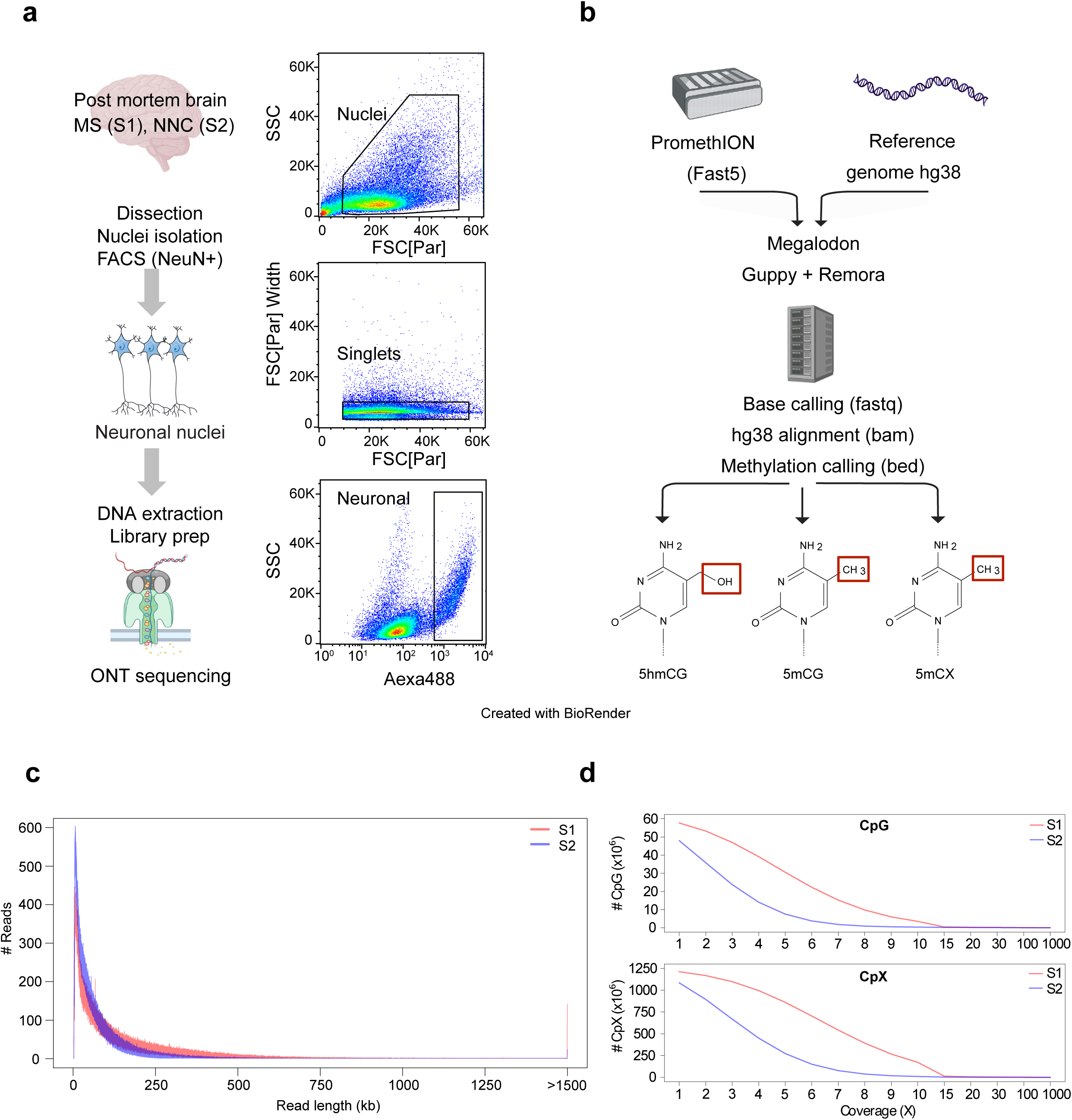
Study design and Oxford Nanopore Technology sequencing. **a)** Post-mortem brain samples were dissected from individuals with Multiple Sclerosis (MS, S1) and non-neurological disease controls (NNC, S2). Isolated nuclei were sorted based on: exclusion of nuclear fragment (top graph), nuclei doublets (middle graph) and NeuN-staining intensity (bottom graph). Genomic DNA was extracted from sorted NeuN+ nuclei, fragmented and profiled with Oxford Nanopore Technology (ONT). **b)** ONT output files were processed through high accuracy base calling algorithms (Guppy) and passed reads were subsequently mapped to the human reference genome, hg38, using Megalodon and minimap2. DNA modification calling in 5(h)mCpG and 5mCpX context was performed with Remora. Line plots illustrating **c)** the read length and **d)** the coverage in CpG and CpX-context of S1 (red) and S2 (blue), respectively.

Thus, we observed an expected long-read distribution covering the vast majority of CpG sites within the human genome.

### Comparison of models for 5hmC and 5mC modification calling

We next sought to compare outputs from the two models, i.e. “CpG” model, which calls 5hmC and 5mC in a CpG context and the “CpX” model, which calls a combined level (5hmC+5mC) in all C contexts. Density plots of DNA methylation levels revealed that separating the 5mC and 5hmC modifications largely altered the distribution. Indeed, the combined 5mC and 5hmC levels called by the “CpX” (5mC+5hmC) model displayed a conventional bimodal distribution with a small fraction of sites displaying methylation around 0%, a large fraction displaying methylation around 100% and median values of 90.9% and 80.8% methylation for S1 and S2 (Supp. Fig. 1a), respectively. However, when only considering the 5mC modification, without 5hmC, using the “CpG” model, DNA methylation levels were largely decreased as reflected in the lower median values of 60% and 58% methylation for S1 and S2, respectively (Supp. Fig. 1a). Noticeably, while the median 5mC levels were similar for S1 and S2, the 5hmC medians differed between samples, with 30% and 13.1% 5hmC in S1 and S2, respectively. Both samples peaked at 0% 5hmC, however, this peak was higher in S2, while S1 displayed a larger shoulder indicating intermediate 5hmC levels (Fig. 2a).

**Figure 2.**
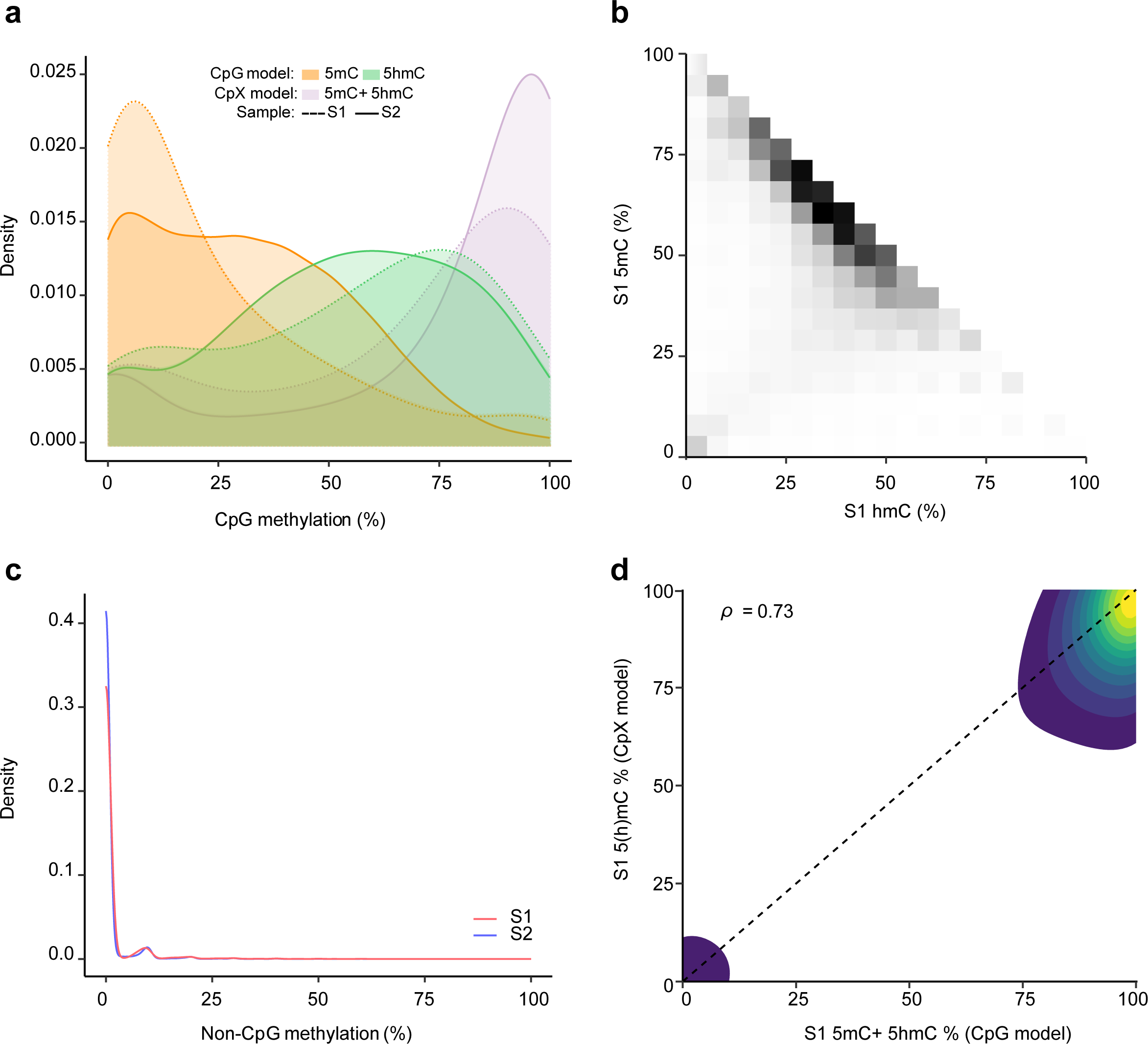
5hmC and 5mC levels in human neurons. **a)** Distribution of 5mC (orange), 5hmC (green), and total 5hmC+5mC (purple) DNA methylation levels (%) in CpG context for S1 (dashed line) and S2 (solid line), respectively. 5hmC and 5mC levels were called with a different algorithm (the CpG model), compared to 5hmC+5mC levels (the CpX model). **b)** Comparison of 5mC and 5hmC levels (%) in 3 kb bins for S1. The white-to-black color gradient illustrates the overlap density. **c)** Density plot of 5mC levels in non-CpG context called with the CpX model for S1 (red) and S2 (blue), respectively. **d)** Comparison of 5hmC+5mC DNA methylation levels called by the CpG (x-axis) and CpX (y-axis) models, respectively. The overlap density is illustrated with a blue-to-yellow colour gradient.

To compare levels of 5hmC and 5mC modifications across genomic locations, we calculated mean 5mC/5hmC levels in 3 kb sliding windows and found that 50-75% 5mC levels most often coincided with 25-50% 5hmC levels (Fig. 2b). High 5mC (100-80%) and low 5hmC (0-10%) were also found, while the opposite scenario of high 5hmC and low 5mC was much less common.

Investigation of 5mC+5hmC levels in a non-CpG context estimated by the “CpX” (5mC+5hmC) model revealed that the majority of non-CpG sites were unmethylated. However, a small proportion of cytosines exhibited DNA methylation levels of ∼10%, as indicated by a peak in the density plot (Fig. 2c). Comparison of methylation levels called by the “CpX” (5mC+5hmC) model, showed strong correlations (ρ_S1_=0.73 and ρ_S2_=0.66) with the 5mC+5hmC levels called by the “CpG” model, although slightly higher levels were called by the former model (Fig. 2d, Suppl. Fig. 1b).

Taken together, we found that while the two models predict comparable DNA methylation levels for 5mC+5hmC in a CpG context, separating the 5hmC and 5mC signal revealed a large contribution of the 5hmC modification.

### Comparison of ONT with array-based BS/oxBS methodology

To benchmark ONT-derived DNA methylation levels against another widely used methodology, we compared ONT with BS/oxBS-based Illumina BeadChip 450K, which enables the detection of both true 5mC and 5hmC. Importantly, both methodologies were applied to the same samples. Comparison of a total of 171,999 sites after applying a 5X filter for ONT and pooling data from both samples yielded Spearman’s correlation coefficients of ρ=0.79 and ρ=0.74 for 5mC and 5hmC, respectively (Fig. 3a, Suppl. Fig. 2). Higher Spearman’s correlation coefficients (on average ρ=0.84 and ρ=0.79 for 5mC and 5hmC, respectively) were obtained with elevated coverage cut-offs of 10X, as exemplified with 14,750 sites in S1 in Fig. 3b (correlation for S2 is shown in Suppl. Fig. 2).

**Figure 3.**
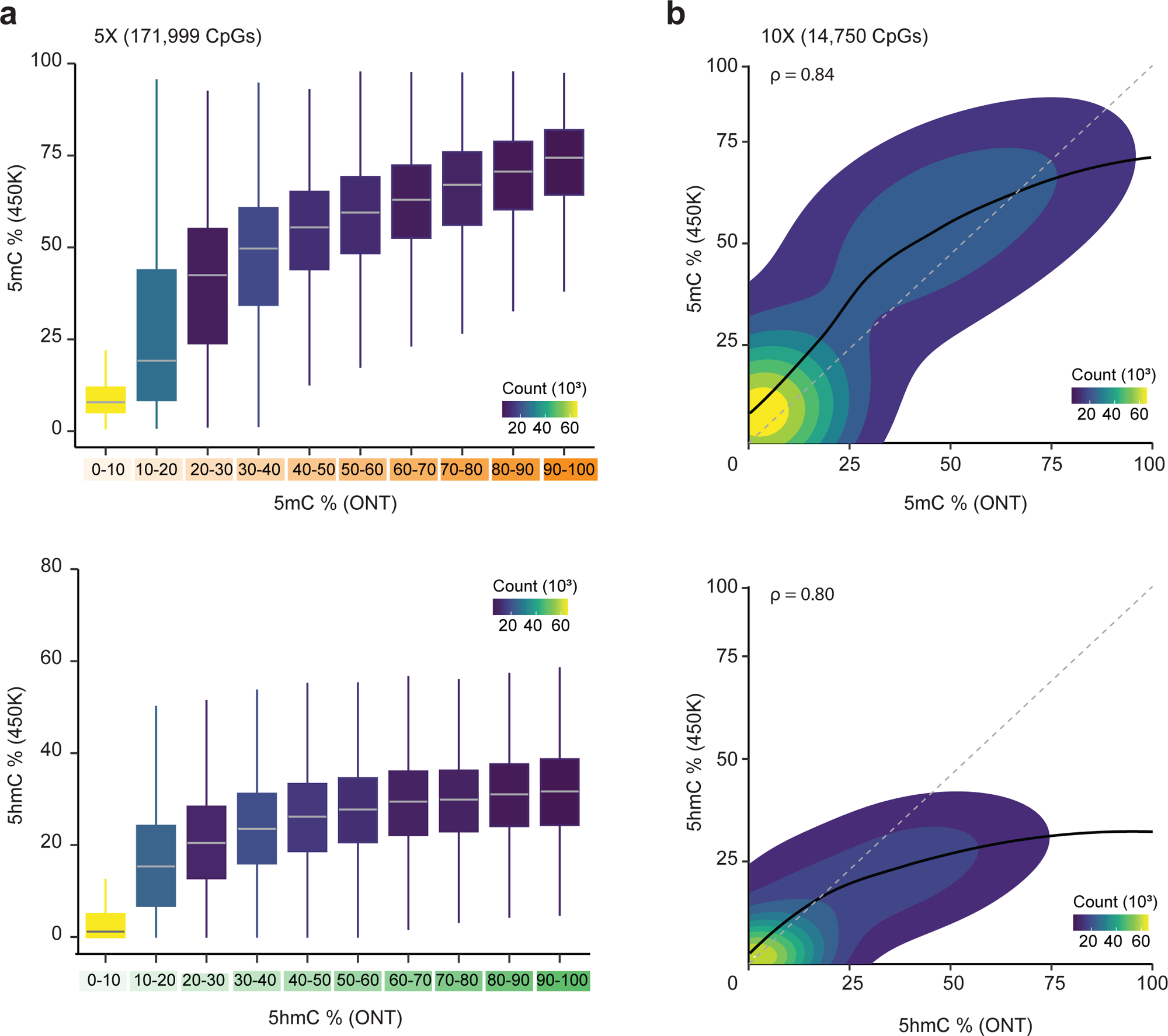
Comparison of ONT and array-based oxBS/BS technology. Comparison of 5mC (upper panel) and 5hmC (lower panel) levels obtained with **a)** ONT (5X coverage, segregated into 10% intervals) and Illumina 450K oxBS/BS methodology. Data from both samples, S1 and S2, was considered. **b)** ONT (10X coverage) and Illumina 450K oxBS/BS methodology for S1. The site number density is depicted in a blue-to-yellow colour gradient.

A “loess” curve of best fit applied to sites passing the 10X ONT cut-off suggested that the relation between 5hmC and 5mC levels from 450K and ONT is not linear (Fig. 3b, Suppl. Fig. 2). This was confirmed by comparing adjusted (adj.) R^2^ values from a linear model with and without polynomial (quadratic and cubic) terms (Table 2, Supp. Figure 2). Indeed, higher adj. R^2^ for a linear model including non-linear polynomial terms suggested that the correlation is not equal across the full range of high, medium and low values.

**Table 2.**
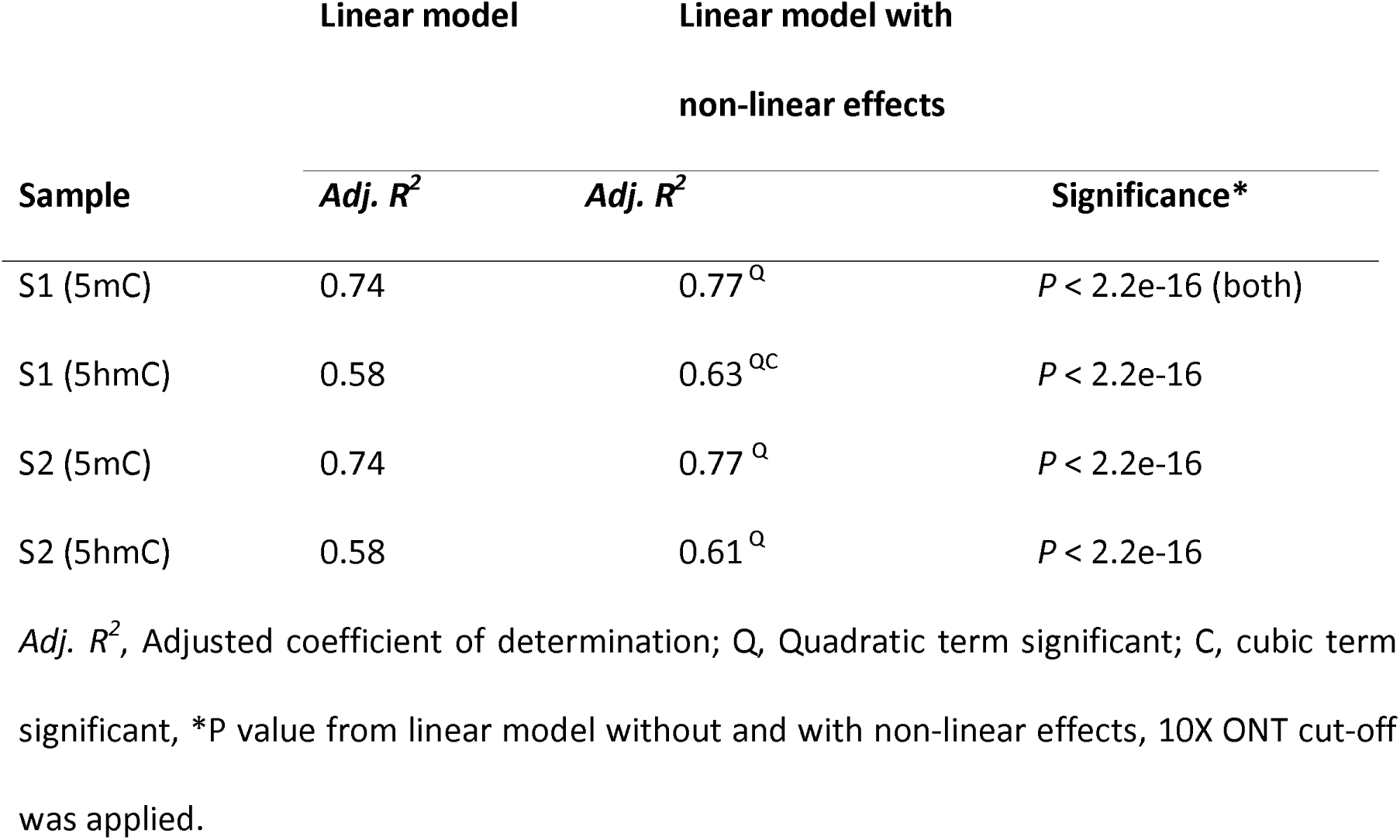
Assessment of non-linear effects (quadratic and cubic) in the correlation between methylation levels from ONT and BS/oxBS (Illumina 450K) methodologies.

### Distribution of neuronal 5hmC and 5mC across genomic features

To explore the genomic distribution of 5hmC compared to 5mC, we examined their average levels across different gene body features. 5hmC levels were generally lower than 5mC levels but followed a similar pattern across the gene (Fig. 4a, Suppl. Fig.3). Intronic CpGs displayed the highest 5mC (50%) and 5hmC (40%) levels, followed by 3’UTR, downstream and coding exon sequences, whereas promoter-like (i.e., upstream and 5’UTR) features exhibited the lowest 5mC (10-45%) and 5hmC (5-30%) levels (Fig. 4a, Suppl. Fig.3). Noticeably, DNA methylation levels of both 5hmC and 5mC decreased at exon-intron junctions (Fig. 4a, Suppl. Fig.3).

**Figure 4.**
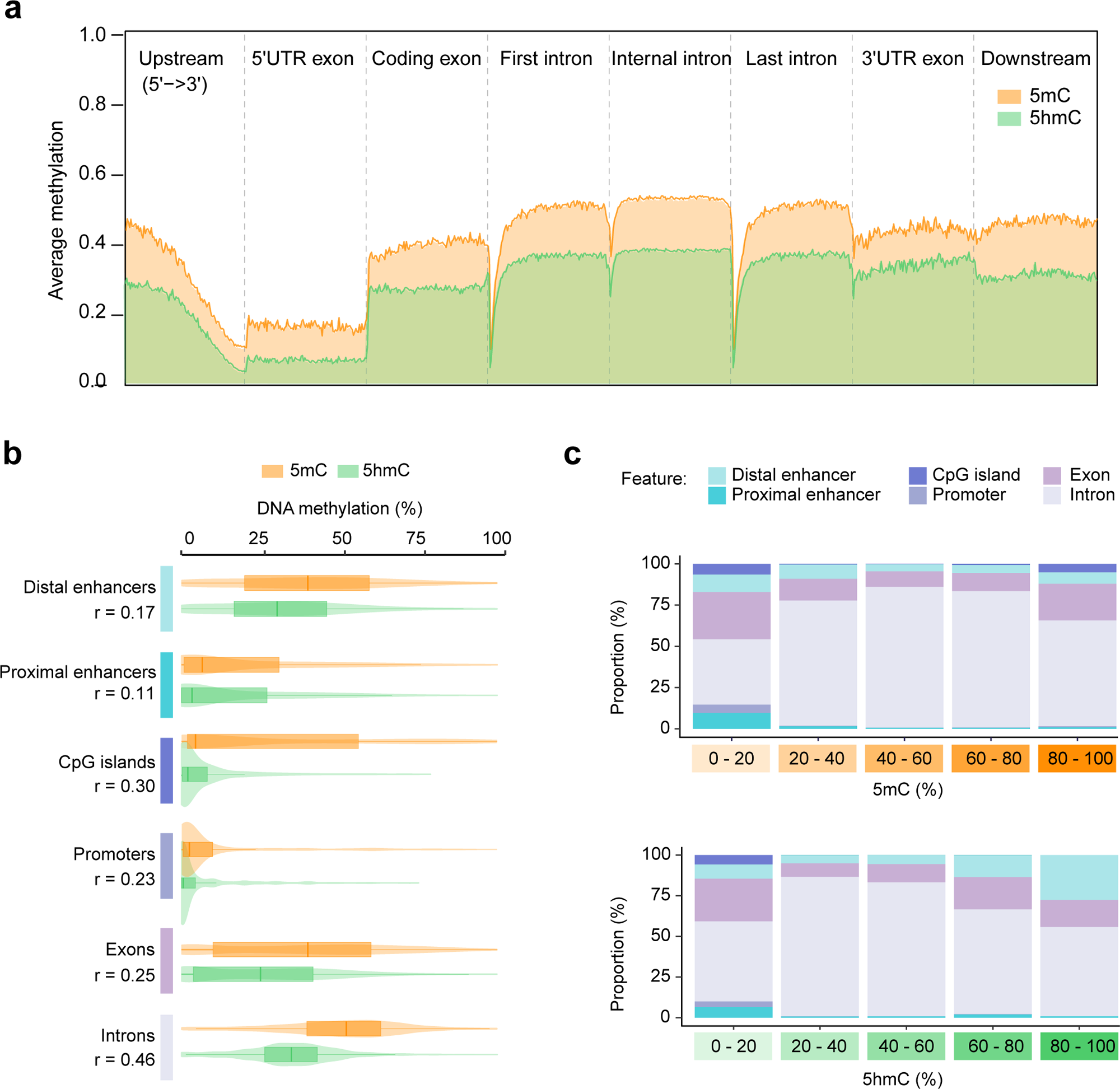
Genomic feature annotation of neuronal 5hmC and 5mC levels. DNA methylation distributions of 5hmC (green) and 5mC (orange) levels (%) was accessed across different **a)** gene- and **b)** regulatory features. UTR: Untranslated region, r: Wilcox effect size **c)** Stacked bar plots illustrating the proportion of different features in segregated 20% bins of 5mC (top) and 5hmC (bottom) DNA methylation levels.

We further explored whether and to what extent 5mC and 5hmC levels differ between generically identified regulatory features such as promoters, proximal enhancers, distal enhancers and CpG islands (CGIs). Comparison of the methylation levels within each feature identified the largest Wilcox effect size differences (indicated with r) between 5mC and 5hmC in introns, with median levels of 52% and 34%, respectively (Fig. 4b). While the distribution of features was similar across low to intermediate 5mC and 5hmC levels, differences emerged at high methylation ranges (Fig. 4c). Indeed, the proportion of distal enhancer elements increased in regions with higher levels of 5hmC, whereas their proportions remained stable across 5mC levels. On the contrary, the proportion of CGIs decreased in regions with higher 5hmC levels, whereas CGIs persisted at the high end of the 5mC methylation range (80–100%) (Fig. 4c).

Thus, while 5hmC and 5mC levels differ, their distribution across regulatory features appears similar at low to intermediate methylation levels, whereas differences are observed in introns, distal enhancers and CGIs in the regions with high methylation levels.

### 5hmC levels are elevated in enhancers and transcribed genes of the human brain

To comprehensively characterize 5hmC and 5mC in neuronal samples, we extended our analysis beyond basic genomic features and focused on characterizing 5hmC and 5mC signals in relation to chromatin states, both within the brain and in the wider contexts of annotations shared across human tissue types. We first explored a universal-stacked ChromHMM model based on IHEC data, where the states were labelled based on a priori universal-stacked model, the updated version including brain-associated enhancers EnhA6 a-d and repressed polycomb ReprPC7 and EnhWk2 features [39]. Hierarchical clustering based on 5hmC and 5mC levels within different states revealed that EnhA6 a+b clustered with EnhWk2, whereas EnhA6 c+d clustered closer to ReprPC7 (highlighted with red text in Fig. 5a). Apart from this, we did not observe apparent differences between the highlighted and other chromatin states in the same enhancer category, suggesting that the universal-stacked ChromHMM model does not directly reflect neuronal 5hmC levels.

**Figure 5.**
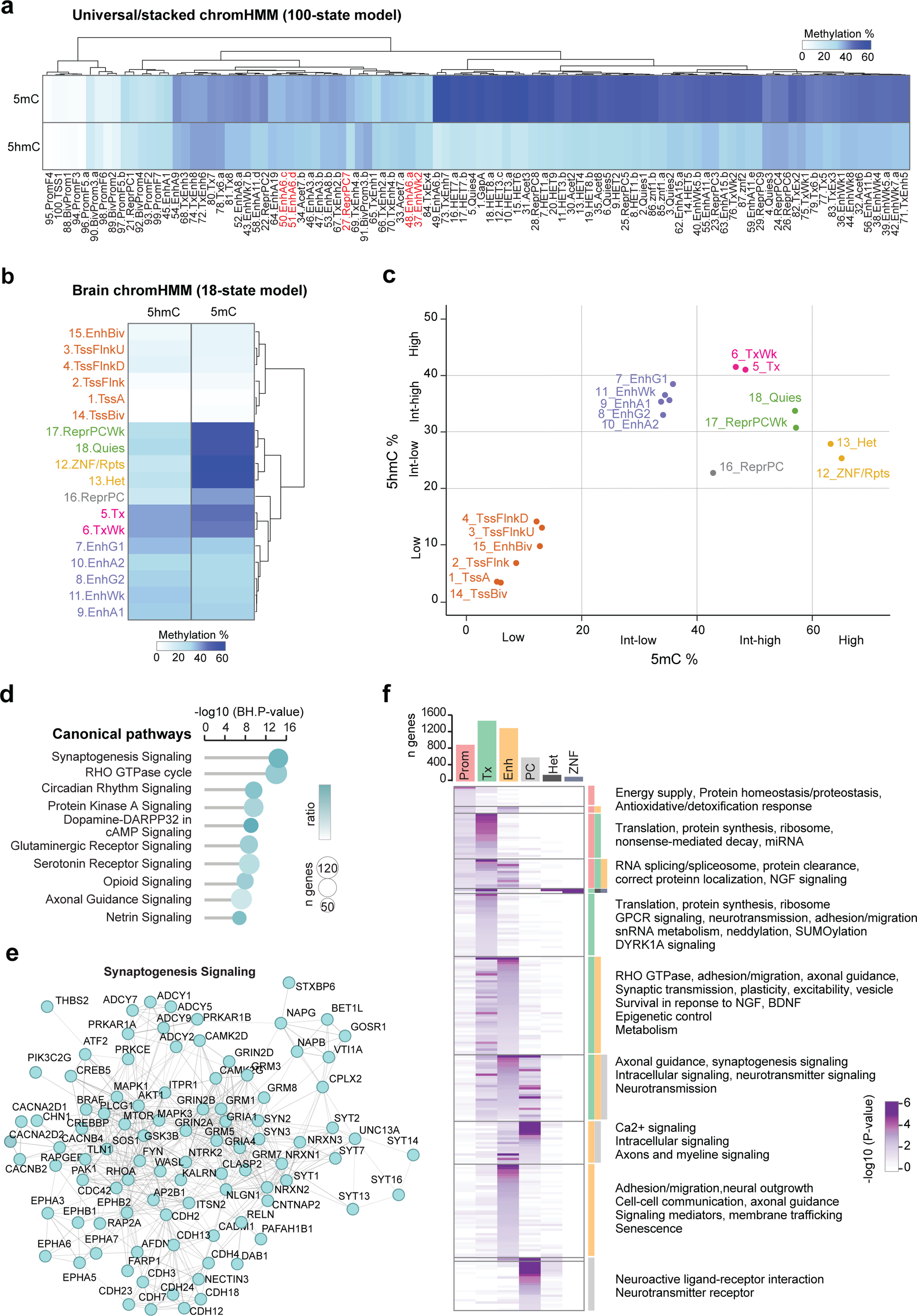
5hmC and 5mC represent different brain-derived chromatin states and have functional relevance for neurons. Heatmaps illustrating average 5hmC and 5mC levels (%) within the **a)** 100-state universal/stacked and **b)** 18-state healthy brain summary chromHMM models. Previously defined brain-specific enhancers are highlighted in red. DNA methylation levels are depicted in a white-to-blue colour gradient. **c)** Scatterplot of average 5hmC (%) and 5mC (%) levels within brain summary chromHMM states color-coded according to their classification (low-intermediate-high). TssA: Active Transcriptional start sites (TSS), TssFlnk: Flanking TSS, TssFlnkU: Flanking TSS upstream, TssFlnkD: Flanking TSS downstream, Tx: Strong transcription, TxWk: Weak transcription, EnhG1: Genic enhancer 1, Enh2: Genic enhancer 2, EnhA1: Active enhancer 1, EnhA2: Active enhancer 2, EnhWk: Weak enhancer, ZNF/Rpts: Zinc finger protein genes and repeats, Het: Heterochromatin, TssBiv: Bivalents/poised TSS, EnhBiv: Bivalent enhancer, ReprPC: Repressed Polycomb, ReprPCWk: Weak repressed Polycomb, Quies: Quiescent/low **d)** Significant canonical pathways predicted from Ingenuity Pathway Analysis (IPA) based on high-5hmC (>60%) input genes. The significance level (-log10(BH-Pval), BH: Benjamini–Hochberg) is depicted in a white-to-turquoise color gradient. The circle size reflects the number of genes (n) represented in each pathway. **e)** Gene network illustration of the top-enriched Synaptogenesis Signaling pathway. **f)** Heatmap illustrating enriched canonical pathways based on high-5hmC (>60%) genes segregated into: Prom: Promoters, Tx: Transcribed, Enh: Enhancers, PC: Polycomb, Het: Heterochromatin and ZNF: ZNF genes. The significance level (-log10(Pval)) is depicted in a white-to-purple color gradient.

Next, we exploited chromatin states summarized across samples classified as “healthy brain” from IHEC. With a representation of more tissue-specific chromatin states, categories separated more clearly according to their 5hmC and 5mC levels (Fig. 5b). Stratification of 5hmC and 5mC levels into 1) low (0-20%), 2) intermediate(I)-low (5hmC: 20-30%, 5mC: 20-40%), 3) intermediate(I)-high (5hmC: 30-40%, 5mC: 40-60%), and 4) high (5hmC: 40-100%, 5mC: 60-100%), further enhanced the separation of chromatin state categories (Fig. 5c). “Low 5hmC & low 5mC” classification represented predominantly states associated with promoter (or TSS) states as well as bivalent enhancers (EnhBiv), “I-high 5hmC & I-low 5mC” represented enhancer states, “high 5hmC & I-high 5mC” included states associated with active transcription (Tx) states. “I-high 5hmC & I-high 5mC” encompassed states representing quiescent/weak polycomb (Quies and ReprPCWk) repressed regions, whereas states repressed by polycomb (ReprPC) classified separately based on “I-low 5hmC & I-high 5mC” levels. Finally, repressed heterochromatin (Het) and zinc finger (ZNF) repeats (Rpts) states associated with “I-low 5hmC & high 5mC” (Fig. 5c).

To gain biological insight into the relevance of 5hmC (Fig. 5c), we performed pathway analysis using Ingenuity Pathways analysis (IPA) of all genes with high (> 60%) 5hmC levels within 2 kb proximity of each chromatin state (n=3,838 genes). Results showed enrichment of canonical pathways associated with neuronal functions, particularly synapse formation, axonal guidance and neurotransmitter signaling, along with general intracellular pathways related to Rho GTPase, protein kinase A and circadian rhythm signaling (Fig. 5d, Suppl. Table 1). The genes implicated in the top term “Synaptogenesis signaling” encode several adhesion molecules of the ephrin, cadherin, neuroligin, neurexin, reelin, synaptotagmin, synapsin, SNAP families together with multiple subunits of glutamate receptors, calcium channels and Rho GTPases, MAPK, cAMP, CREB and downstream signaling (Fig. 5e, Suppl. Table 1). Refinement of the functional annotation according to brain-specific chromatin states implied biological relevance of high 5hmC levels in transcribed (Tx) regions, enhancers and PC repressed segments (Fig. 5f, Suppl. Table 1). More specifically, enriched pathways overlapping with Tx genes were related to common cellular functions such as protein synthesis and splicing, as well as some neuron-related functions such as neurotransmission and axonal guidance. Pathways enriched in genes overlapping with enhancers and polycomb repressed chromatin states seemed neuron-specific and included terms such as Ca^2+^ signaling, axons and myelin signaling, neural outgrowth and neuroactive ligand-receptor interaction (Fig. 5f, Suppl. Table 1).

Altogether, our data imply that 5hmC and 5mC levels are highly associated with brain-derived chromatin states, with high 5hmC levels in transcribed genes, enhancers and polycomb repressed regions with both shared and distinct biological functions.

### Neuronal 5hmC denotes transcriptional regulation

Next, we thought to further explore the regulatory potential of high 5hmC (> 60%) and 5mC (> 80%) regions by conducting transcription factor (TF) motifs enrichment analysis (Fig. 6a). A total of 34 TFs were significantly enriched (adj.Pval < 0.01), when compared to 0% 5hmC (relative analysis, Fig. 6b) with highly significant TFs including Gmeb1, ARNT::HIF1A, TCFL5, E2F8, TFDP1, MAX, E2F6, Ahr::Arnt and Arnt. Noticeably, Gmeb1 and ARNT::HIF1A were also highly enriched in high (> 80%) vs. low (0%) 5mC relative analysis. An additional 13 and 21 TFs passed the adj.Pval < 0.01 threshold for 5hmC and 5mC, respectively, when the absolute enrichment against the background was considered (absolute analysis, Fig. 6b). While nearly 70% of TFs (38 of 55) were significant in regions with both high 5hmC and 5mC, several were specifically associated with either 5hmC, such as E2F8, E2F6 and TFDP1, or 5mC, such as SP1, SP2, SP4 and KLF6, KLF10, KLF12, KLF14 (Fig. 6c). Considering both the relative and absolute analyses for 5hmC and 5mC, the top enriched TFs were Gmeb1, ARNT::HIF1A and ZFP57 (Fig. 6c).

**Figure 6.**
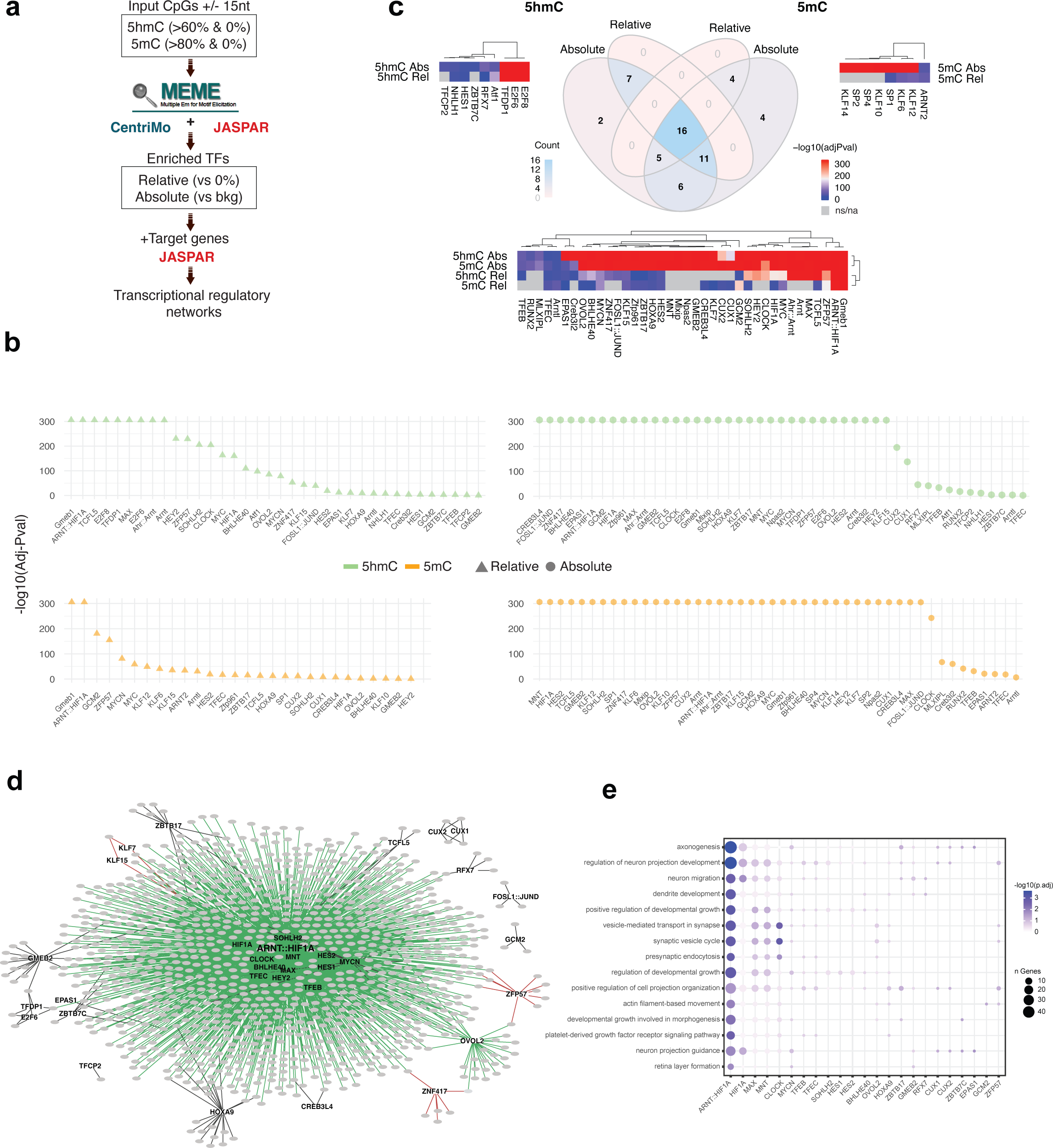
5hmC and 5mC denote transcriptional regulation. **a)** Flowchart illustrating strategy for transcription factor (TF) enrichment and subsequent regulatory network analyses. bkg: background **b)** Scatterplots illustrating significantly enriched TFs (x-axis) ranked according to -log10(adj.Pval). TFs enriched in high-5hmC (green) and high-5mC (orange) sites, above 60% and 80%, respectively, either in comparison to 0% DNA methylated sites (relative, triangle) or against the background (bkg) (absolute, circle) were plotted. **c)** Venn diagram showing overlap between TFs significantly enriched in high-5hmC (60%) and high-5mC (80%) based on absolute or relative approaches, respectively. The color key (blue-to-pink) illustrates the overlap (high-to-low). Heatmaps for high-5hmC (left), high-5mC (right) or overlapping (bottom) TFs. The color key (red-to-blue) illustrates the -log10(adjPval) of the relative (rel) or absolute (abs) enrichment of significantly enriched TFs (high-to-low). **d)** Network of high-5hmC TFs (nodes) and their predicted target genes with high-5hmC (grey circles). Edges colors illustrate whether target genes are significantly enriched (green), depleted (red) or not significant (black) with Fisher’s exact test. **e)** Bobble plot of top-15 GO pathways (rows) with bobble sizes illustrating number of target genes for different TFs (columns). The color key (blue-to-white) illustrates -log10(adjPval).

Given that not only enhancers, but also transcribed genes overlap with 5hmC (Fig. 5c), we next set out to explore transcriptional regulatory networks based on 5hmC-enriched TFs and their 5hmC-high target genes. A dense network formed around ARNT::HIF1A, CLOCK, MAX, and MNT, amongst other TFs (Fig. 6d). GO analysis of network genes showed enriched pathways representing predominantly neuronal and neurodevelopmental related processes, such as axonogenesis, regulation of neuron projection development and neuron migration, with per-TF enrichment confirming the involvement of previously mentioned TFs, especially, ARNT::HIF1A (Fig. 6e). Noticeably, neither Gmeb1 nor ZFP57 were located in the dense part of the network, despite being identified as top-enriched TFs. This was explained by Gmeb1 not existing in JASPAR human motifs, and only 9 (of 98) ZFP57 predicted target genes having high 5hmC levels. Other significantly depleted genes included targets of ZNF417, KLF15 and KLF7. Noticeably, according to the Human Protein Atlas single nuclei data [30], ZFP57 and KLF15 are predominantly expressed in oligodendrocytes and astrocytes, respectively (supp. Figure 4), supporting that their target genes would likely not be expressed in neurons. Taken together, this adds confidence to our approach of exploring transcriptional regulatory networks based on 5hmC-enriched TFs and their targets genes.

## Discussion

In this study, we profiled the whole-genome distribution of 5mC and 5hmC in sorted human neurons using ONT sequencing, compared different DNA methylation calling models, and benchmarked ONT against the array-based BS/oxBS method applied to the same samples. We characterized 5mC and 5hmC in relation to genomic and regulatory features, brain-related and pan-tissue chromatin states, conducted pathway analysis and inferred transcriptional regulatory networks based on high 5hmC regions.

We found that the ONT-based 5mC and 5hmC detection is similar to previously BS/oxBS array-derived quantification [17], even though discrepancies appeared on a base-to-base comparison especially for high 5hmC levels. Moreover, global methylation levels for 5mC, 5hmC (in CpG context) and 5mC (in non-CpG context) also matched previously reported global values from a study investigating methylation levels in sorted mouse neurons using BS/oxBS-sequencing [40].

In our study, the detailed characterization of 5mC and 5hmC across the gene body segments indicated a relative parallelism between 5mC and 5hmC across the gene. Yet, this should be regarded as an average for all genes in neurons and further stratification according to neuronal vs. non-neuronal genes might unveil differences, notably in 5hmC levels, as observed in a study using immunoprecipitation in mouse neurons [5]. Another study exploiting reduced representation oxBS sequencing to investigate 5mC levels in CpG-dense regions in human brain samples concord with our finding [41]. Further, we observed that 5hmC is largely absent from CpG islands (CGIs) and therefore differs from 5mC across this regulatory feature. Low 5hmC in certain CGIs can be supported by results from immunoprecipitation experiments in mouse neuronal progenitor cells, showing that 5hmC is also absent from p300 regions, a CpG-dense regulatory feature [42].

Chromatin states prediction from epigenetic marks with machine learning-based algorithms, such as ChromHMM [43], is a well-established method for assigning tissue-specific functionality. Here we explored recent advancements, including the universal-stacked ChromHMM approach [39], which is based on thousands of epigenome inputs from IHEC [44], and found limited segregation of brain-related enhancers based on their 5hmC and 5mC profiles. Furthermore, we explored chromatin state annotations summarized across samples classified as healthy brain from IHEC and found refined grouping of related ChromHMM states based on 5mC and 5hmC levels. For example, relatively high 5hmC levels were identified across enhancers and states associated with active transcription in the brain tissue [45].

In a study using AbaSI sequencing, the authors explored 5hmC levels in enhancers but came to contradictory findings as they reported elevated 5hmC in poised/bivalent compared to active enhancers [46]. Their findings are based on histone marks from an adult brain midfrontal lobe sample, whereas we used the aforementioned, summarized chromatin state strategy across multiple IHEC brain samples, hence the difference in input material and annotation strategies could potentially have led to different conclusions. Nevertheless, the outcome of pathway-enrichment analysis of 5hmC-enriched genes corroborated the enrichment of terms pertaining to synaptic transmission and neural morphogenesis found by another study using forebrain organoids [47]. Of note, while we show that the combination of 5hmC together with 5mC seem to define brain-related chromatin states, it is important to note that, due to coverage limitations, we did not explore strand-specific DNA methylation levels.

Transcription factor enrichment analysis of highly methylated motifs revealed a substantial overlap in significant TFs between 5hmC and 5mC. Notably, amongst top-enriched TFs - ZFP57 and HIF1A - have previously been shown to bind in a DNA methylation-sensitive manner [48–50]. However, to the best of our knowledge, we are the first to report a significant enrichment in 5hmC across motifs as well. Notably, some of the enriched TF motifs appeared to be specific to 5hmC, including E2F8, E2F6, and their TFDP1 dimerization partner, which are known to repress the cell cycle in post-mitotic cells, such as neurons, through polycomb interaction [51–53], as an important survival mechanism in response to stress and DNA damage [54–57]. In contrast, other distinct classes of TFs - including members of the KLF and SP families - were specifically enriched in 5mC motifs. This is particularly interesting in the light of a study suggesting that KLF4 exhibits differential affinity for various cytosine modifications, potentially allowing a refined response beyond a binary „on/off“ scenario for transcriptional regulation of target genes [58].

Altogether, these findings highlight the continued need for further analysis of 5hmC in a variety of different chromatin states and tissue/cell types to enhance our understanding of the exact mechanisms by which 5hmC shapes cell identity and development.

## Acknowledgements

The computational heavy jobs, such as base- and modification-calling as well as read mapping was enabled by resources provided by the National Academic Infrastructure for Supercomputing in Sweden (NAISS) at UPPMAX, funded by the Swedish Research Council through grant agreement no. 2022-06725. Furthermore, we acknowledge the National Genomics Infrastructure (NGI) at SciLifeLab for ONT sequencing and the support from the Swedish Research Council (VR) and UPPMAX for computational resources. We thank the International Human Epigenome Consortium (IHEC) for providing access to reprocessed and harmonized epigenomic data from a broad collection of human cell and tissue types (https://ihec-epigenomes.org/epiatlas/data/). This study was supported by grants from the Swedish Research Council, the Swedish Association for Persons with Neurological Disabilities, the Swedish Brain Foundation, the Swedish MS Foundation, the Stockholm County Council - ALF project, StratNeuro, Neuroförbundet, the European Research Council grant (grant agreement No 818170) and the Knut and Alice Wallenberg Foundation. L.K. is supported by a fellowship from the Margaretha af Ugglas Foundation.

## Author contributions

DK co-wrote the manuscript, preprocessed and analyzed the ONT dataset and produced the figures 2-4/5. MHS conducted the transcriptional regulatory network analysis and developed the 5hmC_regulator ShinyApp. RAO set up the computational pipeline on the high-performance clusters for base- and modification calling as well as read mapping. HV and JE generated the general tissue and brain ChromHMM annotation for IHEC. LK performed neuronal nuclei isolation, DNA extraction and provided scientific advice, contributed to figures and manuscript. MN provided guiding conceptual, technical and scientific oversight, contributed to analysis, figures and manuscript. MJ provided overarching financial and scientific support, contributed to figures, pathway analysis and the manuscript. All authors read and approved the manuscript.

## Conflict of interest

The authors declare no conflict of interest.

---

**Figure S1. Additional plots for Figure 1 (Methylation overview). a)** Alternative visualisation for Fig. 1a, to represent medians and interquartile ranges for the different models, modifications and samples. **b)** In addition to Fig. 1d, which represents S1 data, we show the same analysis for S2 with lower sequencing depth.

**Figure S2. Comparison Illumina 450k/ONT. a)** Correlation between the methylation levels detected with ONT and 450K for 5X coverage (ONT) sample 2 (n=647 CpGs). **b)** Correlation between the methylation levels detected with ONT and 450K for 10X coverage (ONT) with pooled CpGs from both samples.

**Figure S3. Genomic feature annotation of neuronal 5hmC and 5mC levels in sample 1.** Average DNA methylation distribution of 5hmC (pink) and 5mC (black) levels (%) across gene features. UTR: Untranslated region

**Figure S4. Expression levels of 5hmC-enriched transcription factors.** Heatmap of 5hmC-high enriched transcription factors (columns) across different tissues (rows) available through the human protein atlas. The color key (red-to-white) illustrates expression levels (high-to-low) in count per million (cpm).

## Notes

### Competing Interest Statement

The authors have declared no competing interest.

