## Supplementary figures and images for "Whole-genome profiling of native 5-hydroxymethylation in human neurons with long-read sequencing"

### Supp. Fig. 1

##### Supplementary Figure 1

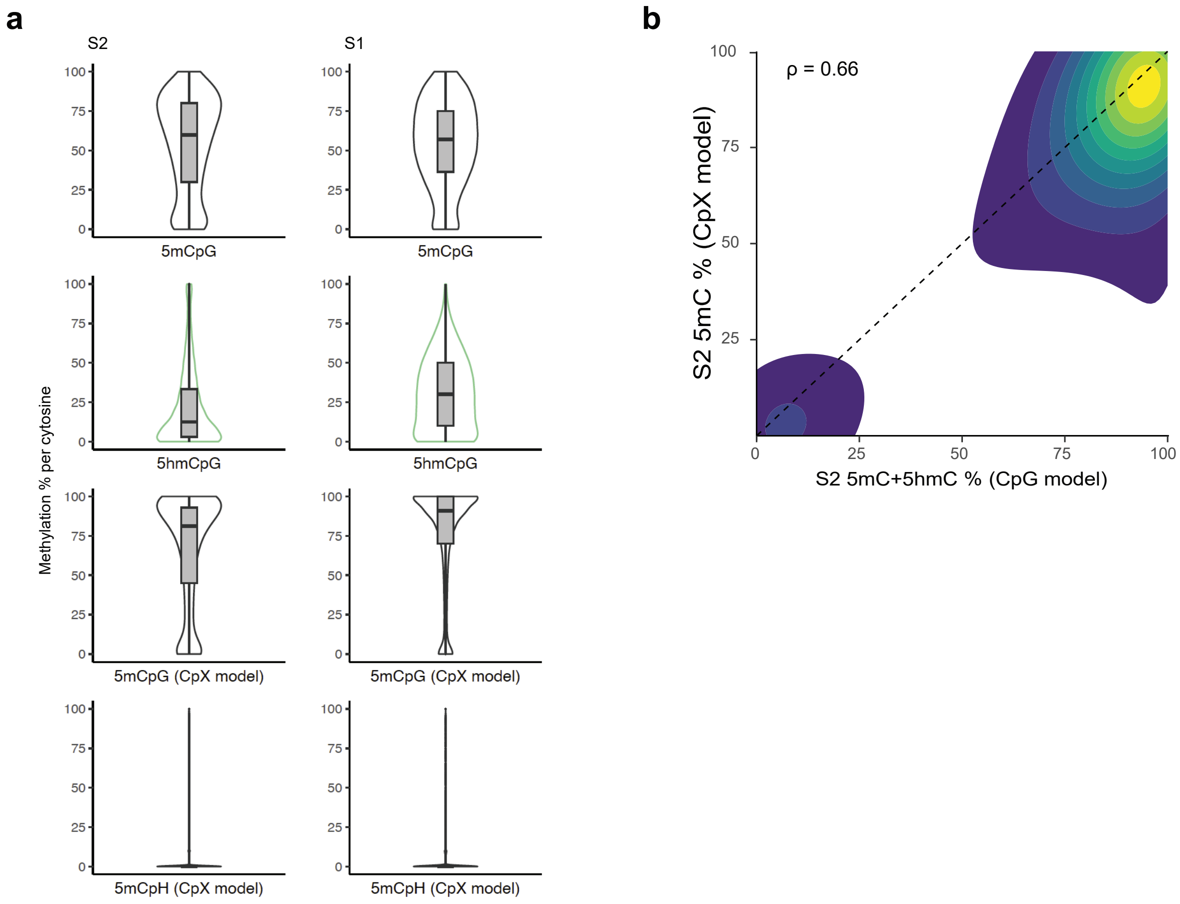

### Suppl. Fig.3

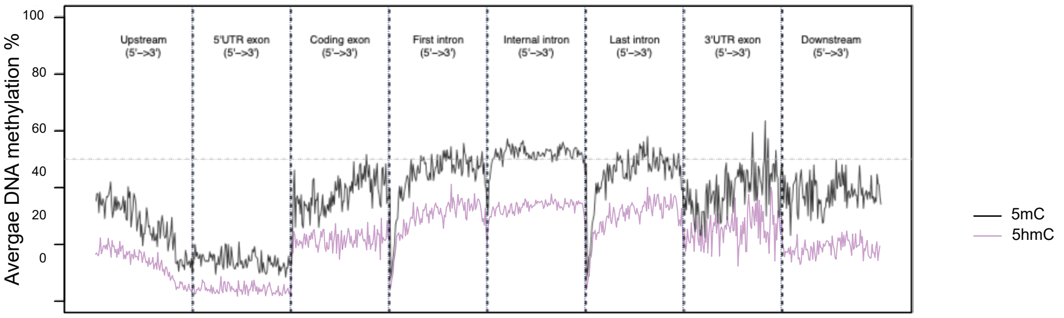

### Suppl. Fig.4

#### Supplementary Figure 4

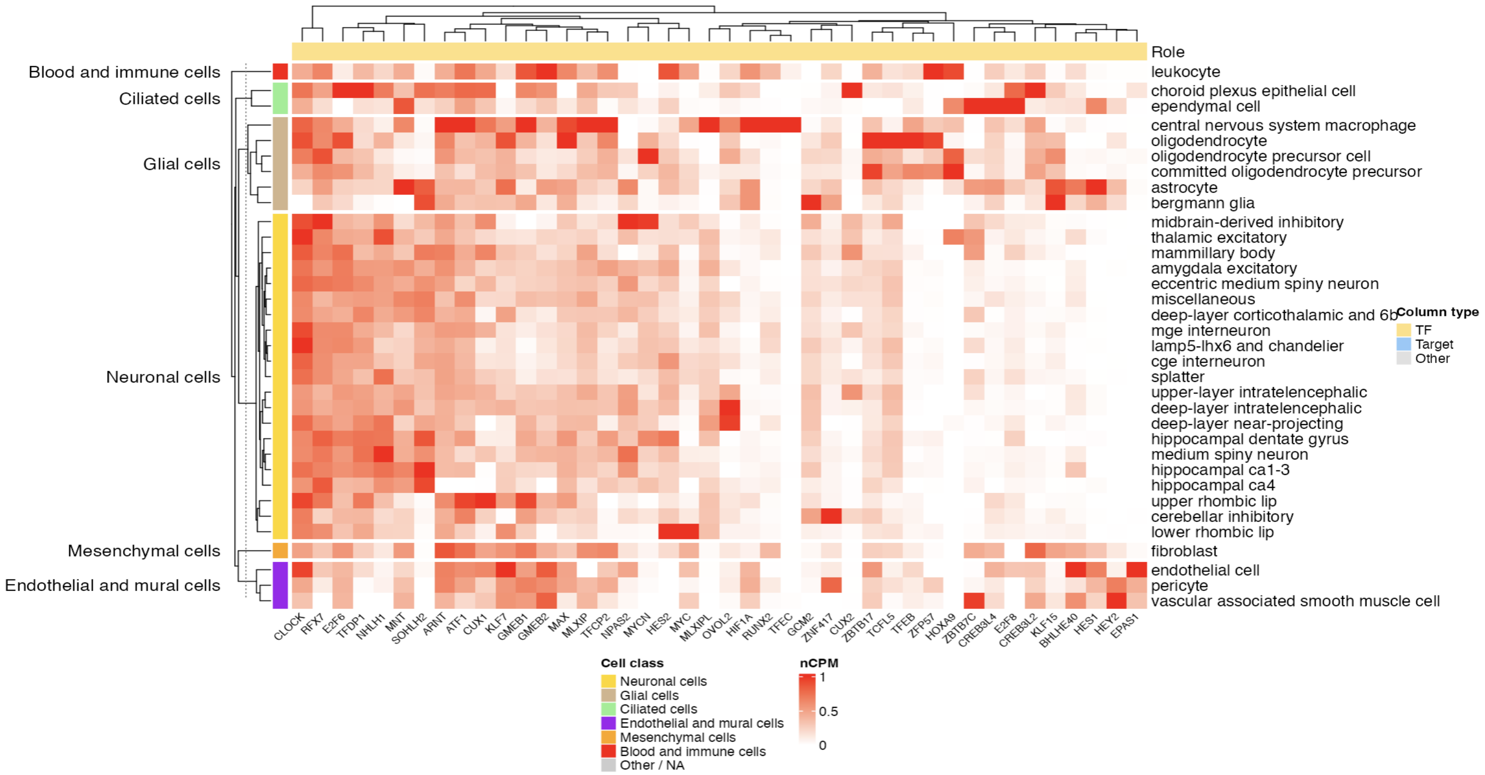

### Suppl. Fig. 2

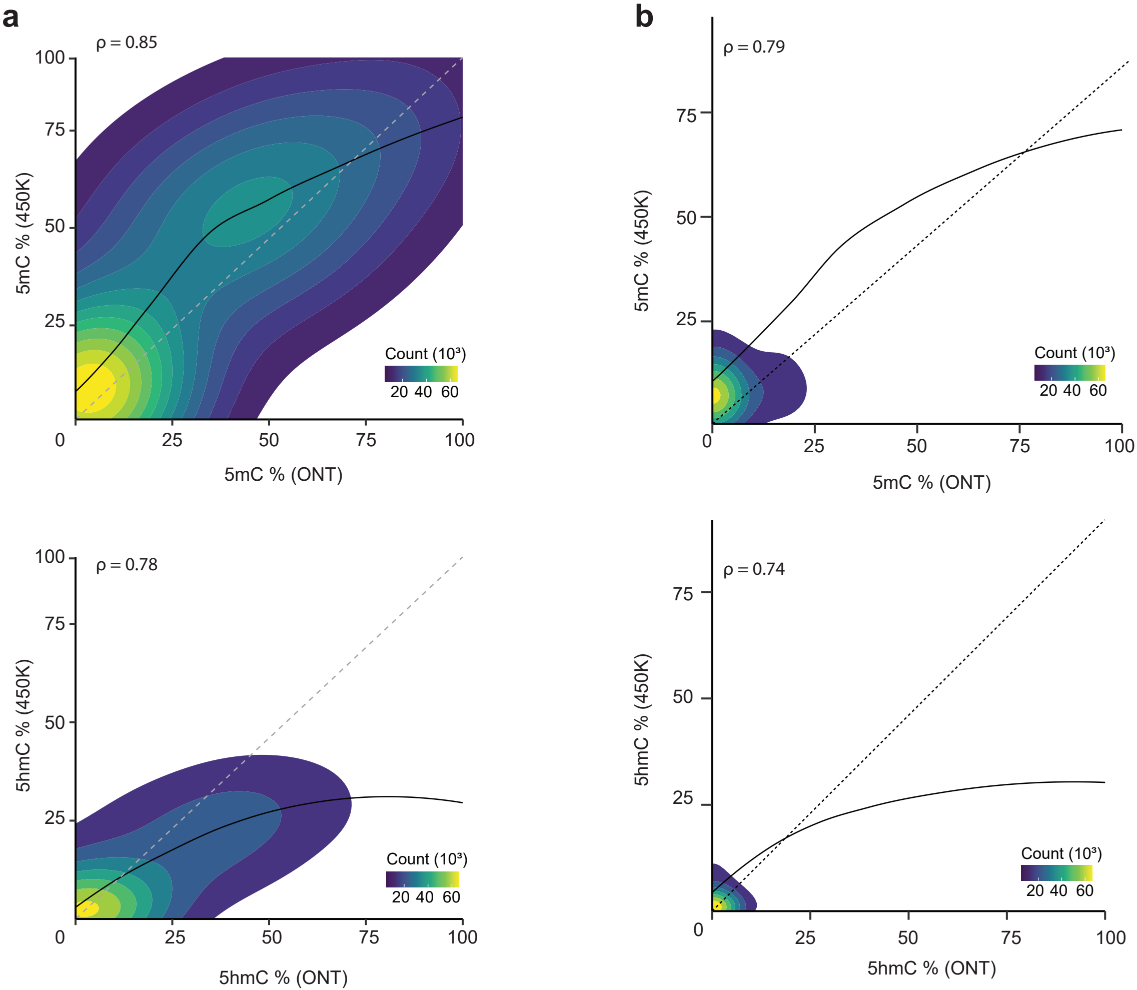
